# On Inferring Additive and Replacing Horizontal Gene Transfers Through Phylogenetic Reconciliation

**DOI:** 10.1101/2020.03.27.010785

**Authors:** Misagh Kordi, Soumya Kundu, Mukul S. Bansal

## Abstract

Horizontal gene transfer is one of the most important mechanisms for microbial evolution and adaptation. It is well known that horizontal gene transfer can be either *additive* or *replacing* depending on whether the transferred gene adds itself as a new gene in the recipient genome or replaces an existing homologous gene. Yet, all existing phylogenetic techniques for the inference of horizontal gene transfer assume either that all transfers are additive or that all transfers are replacing. This limitation not only affects the applicability and accuracy of these methods but also makes it difficult to distinguish between additive and replacing transfers.

Here, we address this important problem by formalizing a phylogenetic reconciliation framework that simultaneously models both additive and replacing transfer events. Specifically, we (1) introduce the *DTRL* reconciliation framework that explicitly models both additive and replacing transfer events, along with gene duplications and losses, (2) prove that the underlying computational problem is NP-hard, (3) perform the first experimental study to assess the impact of replacing transfer events on the accuracy of the traditional DTL reconciliation model (which assumes that all transfers are additive) and demonstrate that traditional DTL reconciliation remains highly robust to the presence of replacing transfers, (4) propose a simple heuristic algorithm for DTRL reconciliation based on classifying transfer events inferred through DTL reconciliation as being replacing or additive, and (5) evaluate the classification accuracy of the heuristic under a range of evolutionary conditions. Thus, this work lays the methodological and algorithmic foundations for estimating DTRL reconciliations and distinguishing between additive and replacing transfers.

An implementation of our heuristic for DTRL reconciliation is freely available open-source as part of the RANGER-DTL software package from https://compbio.engr.uconn.edu/software/ranger-dtl/.

## 1 Introduction

The transfer of genetic information between organisms that are not in a direct ancestor-descendant relationship, called *horizontal gene transfer* or simply *transfer* for short, is a crucial process in microbial evolution. The problem of detecting transfer events has been extensively studied and many different methods have been developed for the problem; see, e.g., [32] for a review. The two most widely used classes of methods are those based on atypical sequence composition and those based on phylogenetic discordance. Sequence composition methods look for atypical dinucleotide frequencies, codon usage biases, or other sequence features that might indicate instances of horizontally acquired genes, but are only effective at short evolutionary time scales and are unable to accurately identify the donors and recipients of transfer events [32], [12]. Phylogenetic methods rely on the fact that horizontal transfers leave tell-tale phylogenetic signatures in the topologies of the transferred genes. These methods construct gene trees for individual gene families and compare them to known species phylogenies to infer possible transfer events. It is well-understood that when a gene is horizontally transferred, it may either add itself as a new gene to the recipient genome, resulting in an *additive* transfer, or replace an existing homologous gene, resulting in a *replacing* transfer [19], [9], [18]. Yet, there do not currently exist any phylogenetic methods that simultaneously model both these types of transfers. This limitation not only affects the applicability and accuracy of these methods but also makes it difficult to distinguish between additive and replacing transfers.

Phylogenetic methods for inferring transfer events can be divided into two classes: (i) Those that implicitly assume that all transfers are replacing transfers and that all discordance between gene trees and species trees is due to these replacing transfer events, e.g., [13], [5], [24], [30], [17], [6], [15], [1], and (ii) those based on the Duplication-Transfer-Loss (DTL) reconciliation framework, which model gene duplication and gene loss as additional sources of gene tree/species tree discordance, but implicitly assume that all transfers are additive transfers, e.g., [11], [23], [31], [10], [8], [2], [27], [28], [29], [25], [16], [20]. Thus, no existing phylogenetic method models both additive and replacing transfers. And while methods based on DTL reconciliation represent a major advance in the ability to accurately detect transfer events, they are limited by their inability to properly handle replacing transfers.

### Our contribution

In this work, we define and formalize a phylogenetic reconciliation framework that simultaneously models both additive and replacing transfer events. Our framework builds upon the standard parsimony-based DTL reconciliation model [31], [2], which assumes that the species tree is undated and seeks an optimal (and not necessarily time-consistent) reconciliation, by explicitly modeling replacing transfer events.^1^ Specifically, we formally define the *<u>D</u>uplication–Additive-Transfer– <u>R</u>eplacing-Transfer– <u>L</u>oss* (DTRL) reconciliation model that explicitly models both additive and replacing transfer events, along with gene duplications and losses. As with the underlying DTL reconciliation model, we formulate the DTRL reconciliation problem as one of finding a *most parsimonious* DTRL reconciliation, i.e., one with smallest total “reconciliation cost”. We prove that the problem of computing a most parsimonious DTRL reconciliation is NP-hard, using a reduction from the NP-hard *minimum rooted Subtree Prune and Regraft (rSPR) distance* problem, and perform the very first experiments to study the impact of replacing transfer events on the accuracy of DTL reconciliation itself. Surprisingly, we found that DTL reconciliation is highly robust to the presence of replacing transfer. Based on these results, we devise a simple heuristic to classify transfer events inferred through DTL reconciliation as being either additive or replacing, and study its classification accuracy using simulated datasets over a range of evolutionary conditions. Our experimental results show that, even though the problem of inferring optimal DTRL reconciliations is NP-hard, it should be possible to design effective heuristics for the problem based on the simpler, and efficiently solvable, DTL reconciliation model.

We note that the problem of integrating replacing transfers with DTL reconciliation has also been recently, and independently, studied by Hasic and Tannier in a recently published manuscript [14]. That manuscript proves that the problem of inferring replacing transfers through phylogenetic reconciliation is NP-hard when the species tree is dated. However, the results in that manuscript are largely complementary to the current work. Specifically, we provide a rigorous and precise formalization of the DTRL reconciliation framework, our proof of NP-hardness is not only completely different but applies to the undated version of the problem where the species tree is undated (arguably the more widely applicable version of the problem), we provide the first experimental results on the impact of replacing transfer on conventional DTL reconciliation, and we devise and evaluate the first heuristic algorithm for estimating optimal DTRL reconciliations.

An abridged version of this paper without proofs appeared in the Proceedings of the 10th ACM International Conference on Bioinformatics, Computational Biology and Health Informatics (ACM-BCB 2019) [21].

The remainder of the manuscript is organized as follows: Basic definitions, preliminaries, and a formal description of the DTRL reconciliation model appear in the next section. The NP-hardness proof appears in Section 3, and experimental results on the effect of replacing transfers on DTL reconciliation are described in Section 4, and our heuristic for classifying transfers is described and tested in Section 5. Concluding remarks appear in Section 6.

## 2 Definitions and preliminaries

We follow basic definitions and notation from [2]. Given a rooted tree *T*, we denote its node, edge, and leaf sets by *V*(*T*), *E*(*T*), and *Le*(*T*) respectively. The root node of *T* is denoted by *rt*(*T*), the parent of a node *v* ∈ *V*(*T*) by *pa*_*T*_ (*v*), its set of children by *Ch*_*T*_ (*v*), and the (maximal) subtree of *T* rooted at *v* by *T*(*v*). The set of *internal nodes* of *T*, denoted *I*(*T*), is defined to be *V*(*T*) \ *Le*(*T*). We define ≤_*T*_ to be the partial order on *V*(*T*) where *x* ≤_*T*_ *y* if *y* is a node on the path between *rt*(*T*) and *x*. The partial order ≥_*T*_ is defined analogously, i.e., *x* ≥_*T*_ *y* if *x* is a node on the path between *rt*(*T*) and *y*. We say that *y* is an *ancestor* of *x*, or that *x* is a *descendant* of *y*, if *x* ≤_*T*_ *y* (note that every node is a descendant as well as ancestor of itself). We say that *x* and *y* are *incomparable* if neither *x* ≤_*T*_ *y* nor *y* ≤_*T*_ *x*. Given a non-empty subset *L* ⊆ *Le*(*T*), we denote by *lca*_*T*_ (*L*) the last common ancestor (LCA) of all the leaves in *L* in tree *T*; that is, *lca*_*T*_ (*L*) is the unique smallest upper bound of *L* under ≤_*T*_. Given *x, y* ∈ *V*(*T*), *x* →_*T*_ *y* denotes the unique path from *x* to *y* in *T*. We denote by *dist*_*T*_ (*x, y*) the number of edges on the path *x* →_*T*_ *y*; note that if *x* = *y* then *dist*_*T*_ (*x, y*) = 0. Given a set *L* ⊆ *Le*(*T*), let *T*′ be the minimal rooted subtree of *T* with leaf set *L*. We define the *leaf induced subtree* of *T* on leaf set *L*, denoted *T* [*L*], to be the tree obtained from *T*′ by successively removing each non-root node of degree two and adjoining its two neighbors. A tree is *binary* if all of its internal nodes have exactly two children. Throughout this work, the *term* tree refers to rooted binary trees.

A *species tree* is a tree that depicts the evolutionary relationships of a set of species. Given a gene family from a set of species, a *gene tree* is a tree that depicts the evolutionary relationships among the sequences encoding only that gene family in the given set of species. Thus, the nodes in a gene tree represent genes. Throughout this work, we denote the gene tree and species tree under consideration by *G* and *S*, respectively. We assume that each leaf of the gene tree is labeled with the species from which that gene (sequence) was obtained. This labeling defines a *leaf-mapping* ℒ_*G,S*_: *Le*(*G*) → *Le*(*S*) that maps a leaf node *g* ∈ *Le*(*G*) to that unique leaf node *s* ∈ *Le*(*S*) which has the same label as *g*. Note that the gene tree can have zero, one, or more than one gene from any species under consideration. The species tree contains at least all the species represented in the gene tree.

### 2.1 Additive and replacing transfers

When a gene is horizontally transferred, there are two possibilities for how it may incorporate itself into the recipient genome. The first possibility is that the transferred gene inserts itself to the recipient genome without overwriting any existing genes, thereby creating a new gene locus for itself. The second possibility is that the transferred gene replaces an existing homologous copy of itself, preserving the total number of genes in the recipient genome; this type of horizontal transfer is sometimes also referred to as xenologous gene displacement [19].

#### Definition 2.1 (Additive transfer).

*An* additive transfer *is a horizontal gene transfer that inserts itself into the recipient genome through the addition of a new gene locus.*

#### Definition 2.2 (Replacing transfer).

*A* replacing transfer *is a horizontal gene transfer that inserts itself into the recipient genome by replacing a homologous gene at an existing gene locus.*

Note that additive transfers result in an increase in the total number of genes in the recipient genome, while replacing transfers do not. We also point out that replacing transfers can only happen if the recipient genome already contains a homologous copy of the gene being transferred. Figure 1 illustrates how additive and replacing transfer events impact the resulting gene tree topology.

**Fig. 1.**
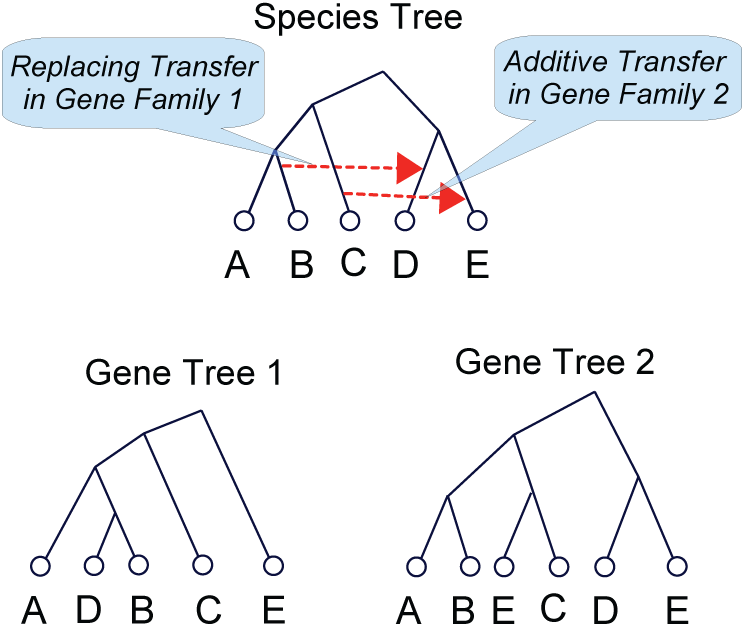
Additive and replacing transfers. This figure shows the evolution of two gene families inside the same species tree. Both gene families exist in the root of the species tree and evolve according to the topology of the species tree without any gene duplications or losses. Gene family 1 is affected by a replacing transfer event, as shown in the figure by the upper orange (dashed) arrow. Gene family 2 is affected by an additive transfer event, as shown by the lower orange (dashed) arrow. The topologies of the resulting gene trees for these two gene families are shown.

### 2.2 DTRL Reconciliation

The *<u>D</u>uplication–Additive-Transfer– <u>R</u>eplacing-Transfer– <u>L</u>oss* (DTRL) Reconciliation model is based upon the well-studied parsimony-based DTL reconciliation framework [31], [2] (which implicitly assumes that all transfer events are additive). However, the introduction of replacing transfers into the model poses several challenges, as we describe below, and the DTL reconciliation framework must therefore be substantially extended to allow for replacing transfers. Specifically, to fully specify a DTRL reconciliation, we must (i) account for hidden duplication or transfer events that do not label any node of the gene tree, and (ii) include in the reconciliation those gene lineages that have been lost (i.e., are no longer visible on the gene tree) but which played a role in the evolution of that gene family by participating in transfer events. We elaborate on these below.

#### Hidden events

Unlike the DTL reconciliation model, where each speciation, duplication, or transfer event required by the reconciliation can be assigned to an individual gene tree node, a most-parsimonious DTRL reconciliation may postulate duplication and transfer events (additive or replacing) that cannot be assigned to any node on the gene tree. Such *hidden events* may be required for most-parsimonious DTRL reconciliation but are invisible on the gene tree either because only descendants from one of the loci resulting from a duplication or additive transfer event survive in the gene family or because they appear on an invisible lineage. The reason hidden events can occur in optimal DTRL reconciliations is that one of the loci resulting from the hidden event is subsequently used (and overwritten) by one or more replacing transfers. This phenomenon is illustrated in Figure 2.

**Fig. 2.**
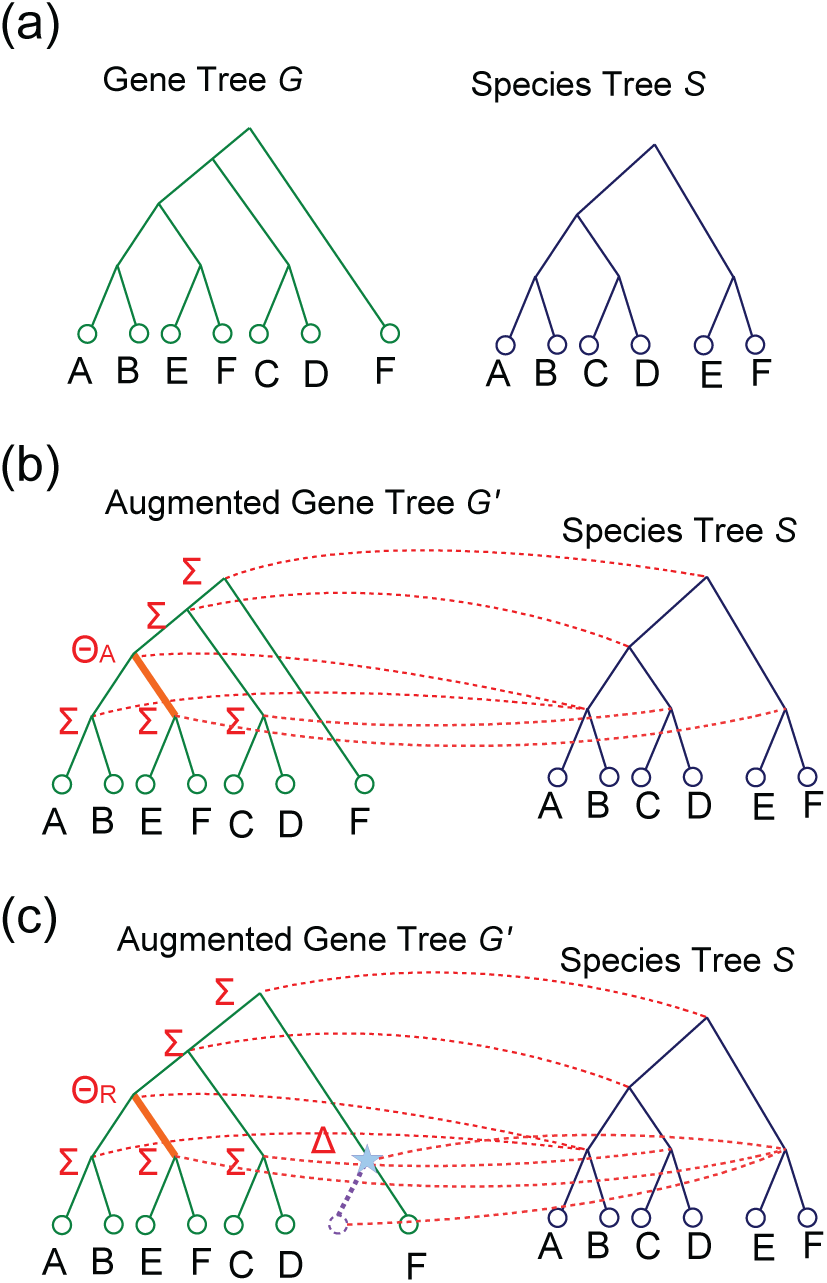
Hidden events and augmented gene trees. Parts (b) and (c) of the figure show two alternative DTRL reconciliations for the gene tree *G* and species tree *S* shown in Part (a). Each reconciliation shows the augmented gene tree *G*′, the event type for each internal node in the augmented gene tree (where Σ represents speciation, Δ represents duplication, Θ_A_ represents additive transfer, and Θ_R_ represents replacing transfer), and the red arcs show the mapping for each node of *G*′ not in *Le*(*G*) (the mapping for each leaf node of *G* is implicitly defined by its leaf label). The bold orange edges represent transfer edges. The reconciliation in Part (b) invokes an additive transfer event and a loss event. For this reconciliation in Part (b), *G*′ is the same as *G*. The reconciliation in Part (c) invokes a replacing transfer event, a hidden gene duplication event (marked by the blue star), and a loss event. The invisible lineage replaced by the replacing transfer event is shown by the purple dotted line in *G*′.

#### Invisible gene lineages and augmented gene trees

To properly recover replacing transfer events and correctly count the number of losses, it is necessary to postulate and account for those gene lineages that are no longer visible on the gene tree but which played a role in the evolution of that gene family by participating in replacing transfer events. Such invisible gene lineages can result from duplication, speciation, or transfer events, but become invisible because no descendants survive in the extant gene family. If these lineages do not participate in any transfer events that impacted the rest of the gene tree, then they can be safely ignored, but otherwise they must be accounted for if replacing transfers are to be recovered accurately and the number of losses counted correctly. We account for invisible lineages by augmenting the input gene tree with additional edges/subtrees, resulting in an *augmented gene tree*, and showing the DTRL reconciliation for this entire augmented gene tree. Figure 2 shows an example of an augmented gene tree and illustrates why it is important to consider invisible gene lineages.

The DTRL reconciliation model takes as input a rooted gene tree and a rooted species tree and defines a framework for reconciling the gene tree with the species tree by postulating duplication, additive transfer, replacing transfer, and gene loss events. The reconciliation creates an augmented gene tree, maps each augmented gene tree node to a unique species tree node, respecting the temporal constraints implied by the species tree topology, and designates each augmented gene tree node as representing either a speciation, duplication, additive transfer, or replacing transfer event. For any gene tree node, say *g*, that represents a transfer event, the reconciliation also specifies which of the two edges (*g, g*′) or (*g, g*″), where *g*′, *g*″ denote the children of *g*, represents the transfer edge and identifies the recipient species of the corresponding transfer. If *g* represents a replacing transfer event, the reconciliation also identifies the specific gene lineage that was lost as a result of that replacing transfer.

Next, we define what constitutes a valid DTRL reconciliation.

##### Definition 2.3 (DTRL-reconciliation).

*A DTRL-reconciliation for G and S is a ten-tuple* ⟨ℒ, *G*′, ℳ, Σ, Δ, Θ_*A*_, Θ_*R*_, Ξ, *τ*, λ⟩, *where* ℒ: *Le*(*G*) → *Le*(*S*) *represents the leaf-mapping from G to S, G*′ *represents the augmented gene tree*, ℳ: *V*(*G*′) → *V*(*S*) *maps each node of G*′ *to a node of S, the sets* Σ, Δ, Θ_*A*_ *and* Θ_*R*_ *partition I*(*G*′) *into speciation, duplication, additive transfer, and replacing transfer nodes, respectively*, Ξ *is a subset of E*(*G*′) *that represents transfer edges (additive or replacing), τ*: Θ_*A*_ ∪Θ_*R*_ → *V*(*S*) *specifies the recipient species for each transfer event, and* λ: Θ_*R*_ → *Le*(*G*′) \ *Le*(*G*) *is an injective function that associates each replacing transfer event with a lost gene in the augmented gene tree, subject to the following constraints:*

##### Augmented gene tree constraint

1. *G* = *G*′[*Le*(*G*)].

##### Mapping constraints

2) *If g* ∈ *Le(G), then* ℳ*(g) =* ℒ*(g).*
3) *If g* ∈ *I*(*G*′) *and g*′ *and g*″ *denote the children of g, then*,
  a. ℳ(*g*) _*S*_ ℳ(*g*′) *and* ℳ(*g*) ≮_*S*_ ℳ(*g*″),
  b. *At least one of* ℳ(*g*′) *and* ℳ(*g*″) *is a descendant of* ℳ(*g*).

##### Event constraints

4) *Given any edge* (*g, g*′) ∈ *E*(*G*′), (*g, g*′) ∈ Ξ *if and only if* ℳ(*g*) *and* ℳ(*g*′) *are incomparable.*
5) *If g* ∈ *I*(*G*′) *and g*′ *and g*″ *denote the children of g, then*,
  a. *g* ∈ Σ *only if* ℳ(*g*) = *lca*(ℳ(*g*′), ℳ(*g*″)) *and* ℳ(*g*′) *and* ℳ(*g*″) *are incomparable*,
  b. *g* ∈ Δ *only if* ℳ(*g*) ≥_*S*_ *lca*(ℳ(*g*′), ℳ(*g*″)),
  c. *g* ∈ Θ_*A*_ ∪ Θ_*R*_ *if and only if either* (*g, g*′) ∈ Ξ *or* (*g, g*″) ∈ Ξ.
  d. *If g* ∈ Θ_*A*_∪Θ_*R*_ *and* (*g, g*′) ∈ Ξ, *then* ℳ(*g*) *and τ*(*g*) *must be incomparable, and* ℳ(*g*′) *must be a descendant of τ*(*g*), *i.e.*, ℳ(*g*′) ≤_*S*_ *τ*(*g*).

##### Replacing transfer constraint

6) *If g* ∈ Θ_*A*_ ∪Θ_*R*_, *then g* ∈ Θ_*R*_ *if and only if* ℳ(λ(*g*)) = *τ*(*g*).

**Note:** This definition allows any invisible leaf node *g* (i.e., *g* ∈ *Le*(*G*′)\ *Le*(*G*)) to map to a leaf node of *S*, say *s* ∈ *Le*(*S*). However, gene *g* is not actually present in species *s* (otherwise it would not be invisible). Instead, ℳ(*g*) = *s* indicates that *g* existed in a predecessor species of *s* represented along the edge (*pa*(*s*), *s*) ∈ *E*(*S*).

In the definition above, Constraint 1 specifies that the augmented gene tree, *G*′, must be consistent with the topology of the input gene tree *G*. Constraint 2 above ensures that the mapping ℳ is consistent with the leaf-mapping ℒ. Constraint 3a imposes on ℳ the temporal constraints implied by *S*, and Constraint 3b implies that any internal node in *G*′ may represent at most one transfer event. Constraint 4 determines the edges of *T* that are transfer edges. Constraints 5a, 5b, and 5c state the conditions under which an internal node of *G*′ may represent a speciation, duplication, and (additive or replacing) transfer respectively. Constraint 5d specifies which species may be designated as the recipient species for any given transfer event. Finally, constraint 6 specifies that a transfer event is labeled as a replacing transfer if and only if there exists a unique invisible leaf node in *G*′ that represents the gene that is “replaced” by that replacing transfer. Note that constraints 2 through 5 are similar to those used in the DTL reconciliation model (e.g. [2]), except that they apply to the augmented gene tree *G*′, not to *G* as in DTL reconciliation, and take replacing transfers into account. Constraints 1 and 6 are unique to DTRL reconciliation.

While duplications, additive transfers, and replacing transfers are directly specified by any DTRL-reconciliation, losses are not. However, given a DTRL-reconciliation, the minimum number of losses implied by that reconciliation can be computed along the same lines as in the DTL reconciliation model [2], but with an adjustment to account for invisible lineages and replacing transfers. The adjustment is required to account for the implicit loss of a gene that occurs at each invisible leaf in the augmented gene tree *G*′. Some of these “losses” are due to replacing transfers, but those that are not must be explicitly counted as gene losses.

###### Definition 2.4 (Losses).

*Given a DTRL-reconciliation α* = ⟨ℒ, *G*′, ℳ, Σ, Δ, Θ_*A*_, Θ_*R*_, Ξ, *τ*, λ⟩ *for G and S, let g* ∈ *I*(*G*′) *and* {*g*′, *g*″} = *Ch*(*g*). *The number of* losses *Loss*_*α*_(*g*) *at node g, is defined to be:*

- *dist*_*S*_(ℳ(*g*), ℳ(*g*′))−1|+|*dist*_*S*_(ℳ(*g*), ℳ(*g*″))−1|, *if g* ∈ Σ.
- *dist*_*S*_(ℳ(*g*), ℳ(*g*′)) + *dist*_*S*_(ℳ(*g*), ℳ(*g*″)), *if g* ∈ Δ.
- *dist*_*S*_(ℳ(*g*), ℳ(*g*″)) + *dist*_*S*_(*τ*(*g*), ℳ(*g*′)) *if* (*g, g*′) ∈ Ξ.

*The number of implicit losses at invisible leaves of G*′ *(i.e., for the set Le*(*G*′) \ *Le*(*G*)*) is defined to be* | *Le*(*G*′) \ *Le*(*G*)| − |Θ_*R*_|.

*The total number of losses in the DTRL-reconciliation α is defined to be Loss*_*α*_ = | *Le*(*G*′)\*Le*(*G*)|−|Θ_*R*_|+∑_*g*∈*I*(*G*)_ *Loss*_*α*_(*g*).

In the DTRL reconciliation framework, each evolutionary event other than speciation is assigned a positive cost. Let *P*_Δ_, 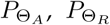, and *P*_*loss*_ denote the gene duplication, additive transfer, replacing transfer, and gene loss costs, respectively. The reconciliation cost of a given DTRL-reconciliation is defined as follows.

###### Definition 2.5 (Reconciliation cost).

*Given a DTRL-reconciliation α* = ⟨ℒ, *G*′, ℳ, Σ, Δ, Θ_*A*_, Θ_*R*_, Ξ, *τ*, λ⟩, *the* reconciliation cost *for α is the total cost of all events invoked by α. In other words, the reconciliation cost of α is* 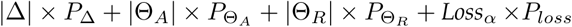.

The goal is to find a DTRL-reconciliation that has minimum reconciliation cost. More formally:

###### Definition 2.6 (ODTRL problem).

*Given G and S, along with P*_Δ_, 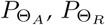, *and P*_*loss*_, *the* Optimal DTRL-Reconciliation Problem (ODTRL) problem *is to find a DTRL-reconciliation for G and S with minimum reconciliation cost.*

## 3 NP-hardness of ODTRL

We claim that the ODTRL problem is NP-hard and that the corresponding decision problem is NP-Complete. The decision version of the ODTRL problem is as follows:

### Problem 1 (D-DTRL).

**Instance:** *G and S, along with event costs P*_Δ_, 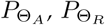, *and P*_*loss*_, *and a non-negative integer l.*

**Question:** *Does there exist a DTRL-reconciliation for G and S with reconciliation cost at most l*

### Theorem 3.1.

The D-DTRL problem is NP-Complete.

The D-DTRL problem is clearly in NP. In the remainder of this section we will show that the D-DTRL problem is NP-hard using a poly-time reduction from the decision version of the NP-hard *minimum rooted Subtree Prune and Regraft (rSPR) Distance* problem [7].

### 3.1 Reduction from minimum rSPR distance

We begin by defining an rSPR operation and define the decision version of the minimum rSPR distance problem.

#### Definition 3.1

(rSPR operation [7]). *Let T be a rooted binary tree and let e* = {*u, v*} *be an edge of T where u is the vertex that is in the path from the root of T to v. Let T*′ *be the rooted binary tree obtained from T by deleting e and then adjoining a new edge f between v and the component C*_*u*_ *that contains u in one of the following two ways:*

- *Creating a new vertex u*′ *which subdivides an edge in C*_*u*_, *and adjoining f between u*′ *and v. Then, either suppressing the degree-two vertex u or, if u is the root of T, deleting u and the edge incident with u, making the other end-vertex of this edge the new root.*
- *Creating a new root vertex u*′ *and a new edge between u*′ *and the original root. Then adjoining f between u*′ *and v and suppressing the degree-two vertex u.*

*We say that T*′ *has been obtained from T by a single* rooted subtree prune and regraft (rSPR) *operation.*

#### Definition 3.2 (rSPR distance).

*Given two trees T and T*′ *with identical leaf sets, the rSPR distance between T and T*′, *denoted d*_*rSPR*_(*T, T*′), *is defined to be the minimum number of rSPR operations required to transform T into T*′.

The minimum rSPR distance problem is to find the rSPR distance between two trees. Its decision version can be stated as follows:

#### Problem 2 (D-rSPR problem).

**Instance:** *Two trees T and T*′ *with identical leaf sets, and a non-negative integer k.*

**Question:** *Is d*_*rSPR*_(*T, T*′) ≤ *k?*

The D-rSPR problem is NP-Complete [7]. Consider any instance *ρ* of the D-rSPR problem with trees *T* and *T*′ on the same leaf set of size *n* (i.e., *Le*(*T*) = *Le*(*T*′) and *n* = | *Le*(*T*)|), and non-negative integer *k*. We will show how to transform *ρ* into an instance *δ* of the D-DTRL problem by constructing *G, S*, and assigning the four event costs *P*_Δ_, 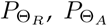, and *P*_*Loss*_, such that there exists a YES answer to the D-rSPR problem on *ρ* if and only if there exists a YES answer to the D-DTRL instance *δ* with reconciliation cost at most *l* = 10*n* + 5*k* − 4.

### 3.2 Gadget

We assume that the leaf set of *T* and *T*′ is {*t*_1_, *t*_2_, …, *t*_*n*_}. We also assume that the internal nodes of *T* are labeled {*z*_1_, *z*_2_, …, *z*_*n*−1_}, as depicted in Figure 3(a). Next, we first show how to construct the species tree *S*, then the gene tree *G*, and then assign event costs.

**Fig. 3.**
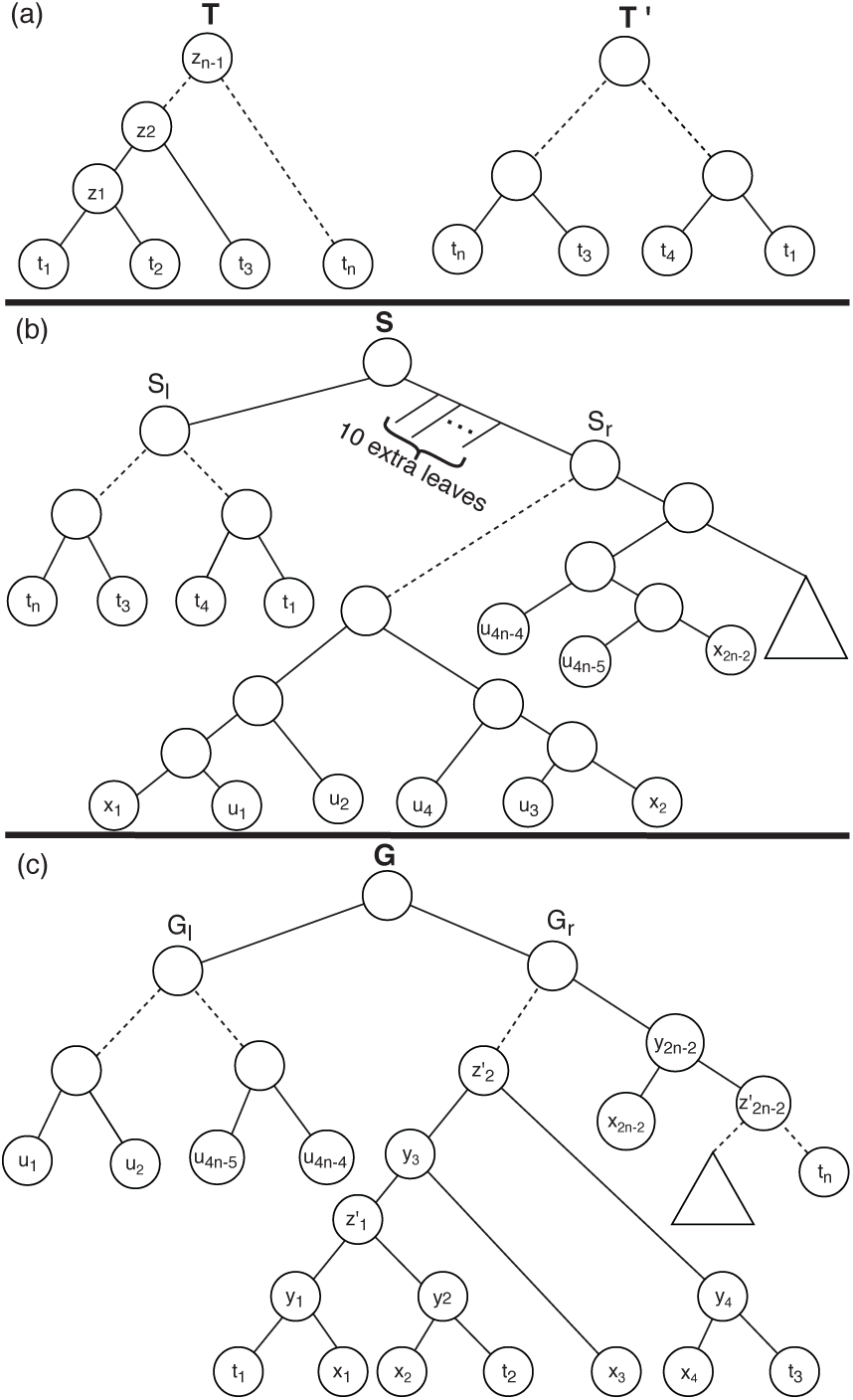
This figure illustrates the construction of species tree *S* (Part (b)) and gene tree *G* (Part (c)) for D-DTRL problem instance *δ* based on trees *T* and *T*′ (Part (a)) in the input instance *ρ* of the D-rSPR problem.

#### Species tree

The species tree *S*, is composed of two subtrees denoted *S*_*l*_ and *S*_*r*_ and ten extraneous leaf nodes (which are not represented in the gene tree). The root of subtree *S*_*l*_ is a child of *rt*(*S*). The other subtree, *S*_*r*_, is connected to *rt*(*S*) through a path to which the ten extraneous leaves are connected; these ten extraneous leaves ensure that no node of *G* maps to *rt*(*S*) in any optimal DTRL reconciliation. This is shown in Figure 3(b). The subtree *S*_*l*_ is identical to tree *T*′. Subtree *S*_*r*_ is a modified version of tree *T*, obtained as follows: We first perform a post-order traversal of tree *T* and number each node according to its position in the ordering, e.g, the left-most leaf node in *T* would be labeled with a 1, while *rt*(*T*) would be assigned the number 2*n* − 1. Next, for each edge (*pa*(*t*), *t*) ∈ *E*(*T*), if the number associated with *t* is *i*, we attach a subtree ((*x*_*i*_, *u*_2*i*−1_), *u*_2*i*_); to edge (*pa*(*t*), *t*). Thus, 2*n*−2 subtrees are attached in all. Finally, we delete all the original leaf nodes {*t*_1_, *t*_2_, …, *t*_*n*_} from *T* and binarize the remaining tree by suppressing all non-root nodes of degree two. The resulting tree is *S*_*r*_. This modification is depicted in Figure 3.

#### Gene tree

Gene tree *G* consists of two main subtrees, denoted *G*_*l*_ and *G*_*r*_. Subtree *G*_*l*_ is obtained from species tree subtree *S*_*r*_ by removing all leaf nodes labeled with prefix *x* and then suppressing all non-root nodes of degree two. Subtree *G*_*r*_ is obtained by modifying *T* as follows: We consider again the post-order numbering of the nodes of *T* and, for each edge (*pa*(*t*), *t*) ∈ *E*(*T*), if the number associated with *t* is *i*, we attach a leaf labeled *x*_*i*_ to edge (*pa*(*t*), *t*). The new internal node created in attaching leaf *x*_*i*_ to the tree is denoted *y*_*i*_. This construction is depicted in Figure 3(c)

Observe that each internal node of *T* has a corresponding node in *G*_*r*_. We label these corresponding nodes of *G*_*r*_ as 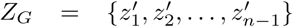, where node 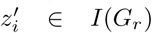 corresponds to node *z*_*i*_ ∈ *I*(*T*) for 1 ≤ *i* ≤ *n* − 1. We also define the following three subsets of *V* (*G*): *Y*_*G*_ = {*y*_1_, *y*_2_, …, *y*_2*n*−2_}, *X*_*G*_ = {*x*_1_, *x*_2_, …, *x*_2*n*−2_}, and *T*_*G*_ = {*t*_1_, …, *t*_*n*_}. Note that *I*(*G*_*r*_) = *Y*_*G*_ ∪ *Z*_*G*_.

**Event costs.** Event costs are assigned as follows: *P*_Δ_ = 4, 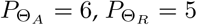, and *P*_*loss*_ = 3.

This completes our construction of instance *δ* of the D-DTRL problem. Note that *G* and *S* can be both constructed in time polynomial in *n* = | *Le*(*T*)|.

**Claim 1.** *There exists a YES answer to the D-rSPR problem on ρ if and only if there exists a YES answer to the D-DTRL instance δ with reconciliation cost l* ≤ 10*n* + 5*k* − 4.

The correctness of Theorem 3.1 follows immediately from Claim 1.

The remainder of this section is devoted to proving this claim, thereby establishing Theorem 3.1. The main idea behind our reduction can be explained briefly as follows. Each rSPR operation on instance *ρ* corresponds to exactly one replacing transfer event on gene tree *G* from instance *δ*. Based on the structure of gene tree *G* and species tree *S*, we will be able to show that for each rSPR operation there is at least one way to get a valid corresponding replacing transfer.

Next, we prove the forward and reverse directions of the claim.

### 3.3 Proof of Claim 1: Forward direction

Assuming we have a YES answer for the *rSPR* instance *ρ*, we will show how to construct a DTRL-reconciliation *α* for instance *δ* with reconciliation cost at most 10*n* + 5*k* − 4.

Suppose *d*_*rSPR*_(*T, T*′) = *k*′, where *k*′ ≤ *k*. Then, based on the close association between rSPR distances and maximum-agreement forests [7], we know that *d*_*rSPR*_(*T, T*′) = *m*(*T, T*′), where *m*(*T, T*′) is the size of a maximum-agreement forest for *T* and *T*′. In particular, there exist *k*′ rooted, vertex-disjoint subtrees of *T*, denoted *T*_1_, …, *T*_*k*′_ with leaf sets ℒ_1_, …, ℒ_*k*′_, respectively, such that *T* [ℒ_*i*_] = *T*′[ℒ_*i*_] for all *i* ∈ {1, …, *k*′}, and ℒ_1_ ∪ … ∪ ℒ_*k*′_ = *Le*(*T*). These *k*′ subtrees from the maximum-agreement forest correspond to the *k*′ subtrees that are pruned and regrafted to transform *T* into *T*′ through rSPR operations. In other words, there exist *k*′ nodes, denoted 𝒫 = {*p*_1_, …, *p*_*k*′_} in *V*(*T*), corresponding to the roots of the *k*′ subtrees *T*_1_, …, *T*_*k*′_, respectively, that identify the edges that will be cut in the *k*′ rSPR operations. For brevity, we refer the reader to [7] for a definition of maximum-agreement forests and for proofs of the preceding statements.

The following observation states three simple facts about the set of nodes 𝒫.

**Observation 1.** *Let t* ∈ *V*(*T*) *and Ch*(*t*) = {*t*′, *t*″}.

1. *If t, t*′ ∈ 𝒫, *then t*″ ∉ 𝒫.
2. *If t*′, *t*″ ∈ 𝒫, *then t* ∉ 𝒫. *Moreover, the set* (𝒫 \ *t*′) ∪ *t must also correspond to a valid maximum-agreement forest for T and T*′.
3. |𝒫| = *k*′ ≤ *k* ≤ *n* − 2.

Parts (1) and (2) in the above observation follow directly from the definition of a maximum-agreement forest. Part (3) follows from the fact that the maximum rSPR distance between any two rooted trees with *n* leaves is bounded above by *n* − 2 [26].

**Notation**: Note that both leaf nodes and internal nodes of *T* have corresponding nodes in the gene tree subtree *G*_*r*_. We denote by 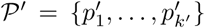 the nodes corresponding to 𝒫 = {*p*_1_, …, *p*_*k*′_} in *G*_*r*_.

Next, we show how to construct the augmented gene tree *G*′ and reconcile *G*′ with *S* such that the total reconciliation cost is no more than 10*n*+5*k*−4. We begin by showing how to reconcile *G* with *S* and then show how to augment *G* into *G*′ and complete the reconciliation. It is worth noting that we start out with 𝒫′ as initialized above, but change its composition as we proceed with defining the reconciliation; however, we will always maintain |𝒫′| = *k*′.

#### Reconciliation of *G* and *S*

We begin by defining a useful edit operation for reconciliations.

##### Definition 3.3 (Switch-recipient operation).

*Given a partial reconciliation of G and S (i.e., a reconciliation in which some nodes of G may not yet have been assigned an event or mapping), and a node g* ∈ *V*(*G*) *that is labeled as a (replacing or additive) transfer event, let g*′ *and g*″ *denote the two children of g such that* (*g, g*′) *is the transfer edge. A* switch-recipient *operation on g, denoted SR*(*g*), *modifies the partial reconciliation by setting* (*g, g*″) *to be the transfer edge, removing edge* (*g, g*′) *from the set* Ξ, *and updating the mappings* ℳ(*g*) *and τ*(*g*) *to be* ℳ(*g*′). *Note that the partial reconciliation of G and S need not remain a valid DTRL-reconciliation after this operation.*

Note: As will become apparent later, the purpose of switch-recipient operations (and also the purpose of the *Y*_*G*_ nodes in the gadget) is to allow for the option of making a transfer node *g* and its sibling have incomparable mappings in *S*. Doing so makes it possible to label the parent of *g* as a transfer as well.

The leaf-to-leaf mapping from *G* to *S* is defined by the leaf labels. To define the remainder of the reconciliation, we first perform a post-order traversal of *G*_*l*_ and map each internal node *a* ∈ *I*(*G*_*l*_) to the species node *lca*(ℳ(*b*), ℳ(*c*)), where *b, c* denote the two children of *a*, and assign *a* to be a speciation event. Next, we perform a post-order traversal of *G*_*r*_ and map each internal node *a* ∈ *I*(*G*_*r*_), where *b* and *c* denote its two children, as described below.

Observe that *I*(*G*_*r*_) = *Y*_*G*_ ∪ *Z*_*G*_, that *X*_*G*_ ∩ 𝒫′ = ø, that every node from *Y*_*G*_ has exactly one child in *X*_*G*_, and that every node from *Z*_*G*_ has both its children from *Y*_*G*_.

1. If *a* ∈ *Y*_*G*_ and *b* ∈ *X*_*G*_ then:
  a. If *c* ∉ 𝒫 ′, then *a* maps to ℳ(*c*) and represents a replacing transfer event with (*a, b*) representing the transfer edge and *τ*(*a*) = ℳ(*b*).
  b. If *c* ∈ 𝒫′, then *a* maps to ℳ(*c*) and represents a replacing transfer event with edge (*a, b*) representing the transfer edge and *τ*(*a*) = ℳ(*b*). We also update 𝒫′ to be (𝒫′ \ {*c*}) ∪ {*a*}.
2. If *a* ∈ *Z*_*G*_ and *b, c* ∈ *Y*_*G*_, then:.
  a. If *a, b, c* ∉ 𝒫′, then *a* maps to *lca*(ℳ(*b*), ℳ(*c*)) and represents a speciation event.
  b. If *a, b* ∉ 𝒫′ and *c* ∈ 𝒫′, then *a* maps to ℳ(*b*) and represents a replacing transfer event and edge (*a, c*) represents the transfer edge with *τ*(*a*) = ℳ(*c*). By Case 1 above, we know that every node of *Y*_*G*_ represents a replacing transfer event, and so *c* must also represent a replacing transfer event. If ℳ(*b*) and ℳ(*c*) are comparable in *S*, i.e., ℳ(*c*) ≤_*S*_ ℳ(*b*) or ℳ(*b*) ≤_*S*_ ℳ(*c*), then we perform the *switch-recipient* operation *SR*(*c*) (which, as we prove later, makes ℳ(*b*) and ℳ(*c*) incomparable).
  c. If *a, c* ∉ 𝒫′ and *b* ∈ 𝒫′, then this case is analogous to the previous case.
  d. If *a* ∉ 𝒫′ and *b, c* ∈ 𝒫′, then
    - If ℳ(*b*) and ℳ(*c*) are incomparable in *S*, then *a* maps to ℳ(*b*) and represents a replacing transfer event with edge (*a, c*) representing the transfer edge and *τ*(*a*) = ℳ(*c*).
    - If ℳ(*b*) and ℳ(*c*) are comparable in *S*, then we perform the *switch-recipient* operation *SR*(*c*). Observe that nodes *b* and *c* must represent replacing transfer events. We also update 𝒫′ to be 𝒫′ = (𝒫′ \ {*b*}) ∪ {*a*}.
  e. If a ∈ 𝒫′ and b, c ∉ 𝒫′, then a maps to lca(ℳ(b), ℳ(c)) and represents a speciation event.
  f. If a, b ∈ 𝒫′ and c ∉ 𝒫′, then a maps to ℳ(c) and represents a replacing transfer with edge (a, b) representing the transfer edge and τ(a) = ℳ(b). If ℳ(b) and ℳ(c) are comparable in S then we perform the switch-recipient operation SR(b) (recall that b must represent a replacing transfer event).
  g. If a, c ∈ 𝒫′ and b ∉ 𝒫′, then this case is analogous to the previous case.
  h. If a, b, c ∈ 𝒫′, then, as we prove later in Lemma 3.1, this case cannot arise in any optimal solution.

Finally, *rt*(*G*) maps to *rt*(*S*_*r*_) and represents an additive transfer event with edge (*rt*(*G*), *rt*(*G*_*r*_)) representing the transfer edge and *τ*(*rt*(*G*)) = ℳ(*rt*(*G*_*r*_)).

Next, we prove some useful properties of the reconciliation described above, show how to augment *G* into *G*′ and “complete” the reconciliation, and prove that the completed DTRL-reconciliation is valid.

##### Lemma 3.1.

*Suppose a* ∈ *Z*_*G*_, *with children b and c, then at no point in the post-order traversal of G*_*r*_, *as described above, can a, b, and c be in the set* 𝒫′ *simultaneously.*

*Proof.* Assume, for contradiction, that *a, b, c* ∈ 𝒫′ at some point during the post-order traversal. Let *a*′ denote the node corresponding to *a* in the tree *T*. Suppose *a*′ ∉ 𝒫. Then, *a* ∉ 𝒫′ at the beginning of the post-order traversal. Observe that *a* cannot be added to 𝒫′ unless the post-order traversal is exactly at node *a* and both *b, c* ∈ 𝒫′ at that time. If *a* is added to 𝒫′ at this step, then one of *b* or *c* will be removed from 𝒫′ and will never be added back at any later time. Thus, if *a*′ ∉ 𝒫 then *a, b, c* ∉ 𝒫′ at any point during the post-order traversal. Consequently, under our assumption, we must have *a*′ ∈ 𝒫.

We will now show that there must exist a node *l* ∈ *Le*(*T*(*a*′)) such that no node along the path from *a*′ to *l*, except for *a*′ itself, is in 𝒫. Consider the two children *u* and *v* of *a*′ in *T*. By part 1 of Observation 1, we know that at most one of *u* or *v* can be in the set 𝒫. Without loss of generality we may therefore assume that *u* ∉ 𝒫. Now, if *u* ∈ *Le*(*T*), then we are done. Therefore, suppose *u* ∉ *Le*(*T*) and let *u*′ and *u*″ denote the two children of *u* in *T*. There are now two possible cases:

1. *v* ∈ 𝒫: In this case, it is not possible that both *u*′ and *u*″ are in the set 𝒫. This is because if *a*′, *v, u*′, *u*″ ∈ 𝒫, then 𝒫\{*a*′}) would yield a valid solution for the D-rSPR problem instance *ρ*, implying *d*_*rSPR*_(*T, T*′) = *k*′ − 1, which is a contradiction.
2. *v* ∉ 𝒫: In this case, if *v* ∈ *Le*(*T*), then we have proved our claim. Therefore, assume *v* ∉ *Le*(*T*) and let *v*′ and *v*″ denote the two children of *v*. Now, it is not possible that *a*′, *u*′, *u*″, *v*′, *v*″ are simultaneously in the set 𝒫. Otherwise, 𝒫\{*a*′}) would yield a valid solution for the D-rSPR problem instance *ρ*, implying *d*_*rSPR*_(*T, T*′) = *k*′ − 1, which is a contradiction.

By applying this argument inductively from *a* towards the leaves of *T*, it follows that there exists a node *l* ∈ *Le*(*T*(*a*′)) such that no node along the path from *a*′ to *l*, except for *a*′ itself, is in 𝒫.

Finally, consider the path from *l* to *a* in *G*_*r*_. This path in *G*_*r*_ consists of nodes corresponding to the *l* to *a*′ path in *T*, along with a subset of nodes from *Y*_*G*_. Observe that, before the post-order traversal of *G*_*r*_, 𝒫′ is initialized to 𝒫 and so none of the nodes along the *l* to *a* path in *G*_*r*_, except for node *a* is in 𝒫′. Furthermore, during the post-order traversal of *G*_*r*_, the current node is added to 𝒫′ only if both children of the current node are in 𝒫′ at that time. Thus, no node along the path from *l* to *a* in *G*_*r*_, except for node *a* can ever be added to the set 𝒫′, and so *a, b*, and *c* cannot simultaneously be in 𝒫′ at any time during the post-order traversal. □

##### Lemma 3.2.

*In the constructed reconciliation of G and S*, ℳ(*z*) ∈ *V*(*S*_*l*_) *for all z* ∈ *Z*_*G*_.

*Proof.* Observe that each node of *T*_*G*_ maps to a node from *S*_*l*_, and that each *z* ∈ *Z*_*G*_ has both children from *Y*_*G*_. To prove that ℳ(*z*) ∈ *V*(*S*_*l*_), for all *z* ∈ *Z*_*G*_, it suffices to prove that, for each *y* ∈ *Y*_*G*_, ℳ(*y*) ∈ *V*(*S*_*l*_) when ℳ(*y*) is first assigned during the post-order traversal of *G*_*r*_. This is because, per case (2) of the post-order traversal, the mapping ℳ(*z*) is assigned based on the initial mapping assignment of the two children of *z*, and while the mapping of one of the children of *z* may be subsequently be changed through a switchrecepient operation, the mapping of *z* remains unchanged.

There are two possible cases:

Case 1: consider any *y* ∈ *Y*_*G*_ such that *y* does not have a child from *Z*_*G*_. In this case, one child of *y* must be in *T*_*G*_ and the other in *X*_*G*_. Since all nodes of *T*_*G*_ map to *S*_*l*_, by case (1) of the post-order traversal we know that the initial mapping assignment for *y* must also be to a node in *S*_*l*_.

Case 2: consider any *y* ∈ *Y*_*G*_ that has a child from *Z*_*G*_. In this case, one child of *y* must be in *Z*_*G*_ and the other in *X*_*G*_. Under a simple inductive argument, we may assume that the child of *y* that is from *Z*_*G*_ maps to a node of *S*_*l*_. Under this assumption, case (1) of the post-order traversal applies and the initial mapping assignment for *y* would therefore be to a node of *S*_*l*_.

A simple inductive argument now immediately establishes that, for each *y* ∈ *Y*_*G*_, ℳ(*y*) ∈ *V*(*S*_*l*_) when ℳ(*y*) is initially assigned during the post-order traversal of *G*_*r*_. □

The next two lemmas helps establish that the assigned transfer events and speciation events are valid.

##### Lemma 3.3.

*In the constructed reconciliation of G and S, if g* ∈ *V*(*G*) *represents a replacing or additive transfer event then* ℳ(*g*) *and τ*(*g*) *must be incomparable.*

*Proof.* Observe that if *g* ∈ Θ_*A*_ ∪Θ_*R*_, then *g* ∈ {*rt*(*G*)} ∪ *Y*_*G*_ ∪ *Z*_*G*_. We therefore have the following three cases:

1. *g* = *rt*(*G*). In this case, based on the constructed reconciliation, *τ*(*g*) = ℳ(*rt*(*G*_*r*_)) and ℳ(*g*) = ℳ(*rt*(*G*_*l*_)). Note that *rt*(*G*_*r*_) ∈ *Z*_*G*_ and so, by Lemma 3.2, *rt*(*G*_*r*_) must map to a node in *V*(*S*_*l*_). Similarly, based on the constructed reconciliation, *rt*(*G*_*l*_) and *rt*(*G*) both map to *S*_*r*_. Thus, ℳ(*rt*(*G*)) and *τ*(*g*) are incomparable.
2. *g* ∈ *Y*_*G*_. Let *g*′ and *g*″ denote the two children of *g*. We know that *g*′ ∈ *Z*_*G*_ ∪ *T*_*G*_ and *g*″ ∈ *X*_*G*_. We know that all nodes of *X*_*G*_ map to nodes of *S*_*r*_, all nodes of *T*_*G*_ map to nodes of *S*_*l*_, and, by Lemma 3.2, all nodes of *Z*_*G*_ map to nodes of *S*_*l*_. Thus, *g*′ must map to a node of *S*_*l*_ and *g*″ must map to a node of *S*_*r*_. Thus, in the initial mapping assignment of *g*, ℳ(*g*) ∈ *V*(*S*_*l*_) while *τ*(*g*) ∈ *V*(*S*_*r*_). Later, if a switch-recipient operation is performed on *g*, we would get ℳ(*g*) ∈ *V*(*S*_*r*_) while *τ*(*g*) ∈ *V*(*S*_*l*_). In either case, ℳ(*g*) and *τ*(*g*) are incomparable.
3. *g* ∈ *Z*_*G*_. Let *g*′ and *g*″ denote the two children of *g*. We know that *g*′ and *g*″ are both in *Y*_*G*_. Based on the case above, we know that each node from *Y*_*G*_ has one child mapping to *S*_*l*_ and the other child mapping to *S*_*r*_. According to case 2 of the post-order traversal, if ℳ(*g*′) and ℳ(*g*″) are comparable (so both map to either *S*_*l*_ or both to *S*_*r*_) then a switch-recipient operation is performed on one of the children of *g*, say *g*′, which would change the mapping of *g*′ from either *S*_*l*_ to *S*_*r*_ or vice versa. Thus, *g*′ and *g*″ are either incomparable to begin with or are made incomparable through a switch-recipient operation. Finally, the mapping of ℳ(*g*) is assigned to be the mapping of one of *g*′ or *g*″, with *τ*(*g*) assigned to be the mapping of the other child. ℳ(*g*) and *τ*(*g*) must therefore be incomparable. □

For the next lemma we need the following definition.

##### Definition 3.4 (*Base Leaf Set*).

*Given the reconciliation of G and S as defined earlier, along with the set* 𝒫′, *we define the* base leaf set *of a node g* ∈ *V*(*G*) *in G, denoted BLe*_*G*_(*g*), *to be* {*l* ∈ *Le*(*G*(*g*)) | *none of the nodes, except possibly g, on the path from g to l is in* 𝒫′}. *We also define BLe*_*S*_(*g*), *for g* ∈ *V*(*G*), *to denote the corresponding set of leaf nodes from S.*

Note that, based on the proof of Lemma 3.1, it follows that | *BLe*_*G*_(*g*)| ≥ 1 for any *g* ∈ *V*(*G*).

##### Lemma 3.4.

*In the constructed reconciliation of G and S, if g* ∈ *I*(*G*) *represents a speciation event and g*′, *g*″ *denote the two children of g, then* ℳ(*g*′) *and* ℳ(*g*″) *must be incomparable in S.*

*Proof.* Based on the constructed reconciliation, if *g* ∈ *I*(*G*) represents a speciation event then either *g* ∈ *I*(*G*_*l*_) or *g* ∈ *Z*_*G*_. We consider these two cases separately.
1. *g* ∈ *I*(*G*_*l*_). In this case, ℳ(*g*) maps to *lca*(ℳ(*g*′), ℳ(*g*″)), and, based on the topologies of *G* and *S*, ℳ(*g*′) and ℳ(*g*″) must be siblings in *S*. Thus, ℳ(*g*′) and ℳ(*g*″) must be incomparable in *S*.
2. *g* ∈ *I*(*G*_*r*_). In this case, based on cases 2(a) and 2(e) of the post-order traversal, we must have *g* ∈ *Z*_*G*_, *g*′, *g*″ ∉ 𝒫′ and *g*′, *g*″ ∈ *Y*_*G*_. Now, observe that for any node *y* ∈ *Y*_*G*_, where *y* ∉ 𝒫′, if *y*′ is the child of *y* that is from *Z*_*G*_ ∪ *T*_*G*_, then *y*′ ∉ 𝒫′ and ℳ(*y*) = ℳ(*y*′). Also observe that if a node *z* ∈ *Z*_*G*_ is not in 𝒫′, then it follows from the proof of Lemma 3.1 that at most one of its two children, denoted *y*′, *y*″, can be in 𝒫′. Furthermore, if *y*′ ∈ 𝒫′, then ℳ(*z*) = ℳ(*y*″), while if *y*′, *y*″ ∉ 𝒫′, then ℳ(*z*) = *lca*(ℳ(*y*′), ℳ(*y*″)). Continuing in this fashion towards the leaves of *G*, it follows that ℳ(*g*′) = *lca*(*BLe*_*S*_(*g*′)) and ℳ(*g*″) = *lca*(*BLe*_*S*_(*g*″)). Since *g* is a speciation node it also follows that *BLe*_*G*_(*g*) = *BLe*_*G*_(*g*′) ∪ *BLe*_*G*_(*g*″) and ℳ(*g*) = *lca*(ℳ(*g*′), ℳ(*g*″)). Consider the induced subtrees *G*[*BLe*_*G*_(*g*)], *G*[*BLe*_*G*_(*g*′)], and *G*[*BLe*_*G*_(*g*″)]. Since none of the edges in these induced subtrees is in 𝒫′, these subtrees must be isomorphic to the induced subtrees *S*[*BLe*_*S*_(*g*)], *S*[*BLe*_*S*_(*g*′)], and *S*[*BLe*_*S*_(*g*″)], respectively. Thus, since *G*(*g*′) and *G*(*g*″) are disjoint subtrees, so must *S*(ℳ(*g*′) and *S*(ℳ(*g*″)), completing the proof □

##### Lemma 3.5.

*In the constructed reconciliation of G and S, there is at most one gene copy in each node (or edge) of S.*

*Proof.* This follows directly from the fact that all internal nodes in *V*(*G*) \ {*rt*(*G*)} represent either speciation or replacing transfer events.

□

We now show how to create the augmented tree *G*′ based on gene tree *G* and construct a complete DTRL-reconciliation. We begin by initializing *G*′ to be the same as *G*,with each node of *G*′ having the same event and mapping assignment as in the reconciliation of *G*. We then perform a post-order traversal of *G*′ and for each node that represents a replacing transfer event, say *g*, we will augment *G*′ by adding a new leaf node, denoted *ū*, connected to *G*′ through a new internal node denoted *<u>u</u>*. This augmentation happens through the *Add*_*G*′_ operation defined below.

##### Definition 3.5

(*Add* operation). *Given G*′, *S, and a node g* ∈ *V*(*G*′) *that is a replacing transfer event, let g*′ *and g*″ *denote the two children of g such that* (*g, g*′) *is the transfer edge and s*′ = ℳ(*g*′). *Note that s*′ ∈ *V*(*S*) \ {*rt*(*S*)} *and so it must have a sibling, which we denote by s*″. *Let u* ∈ *V*(*G*′) *be a node such that* ℳ(*u*) ∈ *V*(*S*(*s*″)), ℳ(*pa*(*u*)) >_*S*_ *s*″ *and* ℳ(*u*) *has minimum distance to the node s*″ *among all options for u. The operation Add*_*G*′_ (*g*) *modifies G*′ *by (i) adding a new node <u>u</u> subdividing the edge* (*u, pa*(*u*)) *(or as new root of G*′ *in the case that rt*(*G*′) = *u), (ii) adding an edge connecting <u>u</u> to a new leaf node denoted* 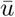, *(iii) assigning to <u>u</u> a <u>m</u>apping of pa*(*s*′) *and event type speciation, and (iv) assigning to* (*u*) *a mapping of s*′.

##### Lemma 3.6.

*For any g* ∈ *G*′ *where g* ∈ Θ_*R*_, *the operation Add*_*G*′_ (*g*) *can be successfully applied.*

*Proof.* Suppose *g* has children *g*′ and *g*″, with (*g, g*′) ∈ Ξ. Based on the constructed reconciliation, if node *g* ∈ Θ_*R*_, then 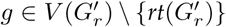. Consequently, ℳ(*g*′) ∈ *V*(*S*_*l*_) ∪ *V*(*S*_*r*_) \ {*rt*(*S*_*l*_)} \ {*rt*(*S*_*r*_}. Thus, if *s*′ = ℳ(*g*′), then *s*′ must have a sibling, say *s*″.

Now, since each leaf *y* ∈ *Le*(*S*_*r*_) ∪ *Le*(*S*_*l*_), has a mapping from a node in *G*′, there must be at least one node *u* that maps to a node in *V*(*S*(*s*″)) and for which ℳ(*pa*(*u*)) >_*S*_ *s*″. Thus, *Add*_*G*′_ (*g*) can be successfully applied. □

##### Lemma 3.7.

*The final augmented gene tree G*′ *is a valid DTRL-reconciliation.*

*Proof.* From Lemmas 3.1 through 3.4 we know that the mapping and event assignments on *G* were valid, and from Lemma 3.6 we know that each *Add* operation itself can be successfully applied. To show that *G*′ is a valid DTRL-reconciliation it therefore suffices to establish the following: (i) the new internal nodes added through the *Add* operations have valid mapping and event assignments, (ii) the parent of each newly added internal node continues to have a valid event and mapping assignment (carried over from *G*), and (iii) each replacing transfer event on *G*′ is associated with a unique lost gene on *G*′.

Consider any new internal node *<u>u</u>* added to *G* through a *Add* operation. By the definition of an *Add* operation, if *u* maps to node *s* in the *S*, then one child of *u* maps to a node from *V*(*S*(*s*″)), and the other child of *u* maps to node *s*′, where *s*′ and *s*″ denote the two children of *s*. Thus, both the mapping assignment and event assignment (of speciation) for *u* are valid.

Now, consider the edge (*v, u*) ∈ *E*(*G*′), where *v* = *pa*(*u*), on which a new internal node *<u>u</u>* is added through an *Add* operation. Let *s* = ℳ(*u*), *s*′, *s*″ ∈ *Ch*(*s*), and, consistent with the definition of an *Add* operation, ℳ(*u*) ∈ *V*(*S*(*s*″)). Observe that, since ℳ(*u*) < ℳ(*v*), *v* could only have been a speciation node in *G*. Moreover, from the definition of an *Add* operation we know that ℳ(*v*) ≥_*S*_ *s*. However, node *v* could not map to *s*, since then the sibling of *u* in *G*′ would map to a node from *V*(*S*(*s*′). But then, *s*′ could not have been the recipient of a replacing transfer event, a contradiction. Thus, ℳ(*v*) >_*S*_ *s*, and so *v* remains a valid speciation event in *G*′ with a valid mapping.

Finally, since an *Add* operation is performed for each replacing transfer node *g* in *V*(*G*) and *Add*_*G*′_ (*g*) adds a corresponding lost gene copy to the gene tree, each replacing transfer event on *G*′ is associated with a unique lost gene on *G*′. □

##### Lemma 3.8.

*If G*′ *denotes the final augmented gene tree, the constructed reconciliation of G*′ *and S does not have any gene losses.*

*Proof.* From Lemma 3.5 we know that each node (edge) on *S* has at most one gene copy. We also know that each leaf node of the species tree node *a* ∈ *Le*(*S*(*S*_*r*_)) ∪ *Le*(*S*(*S*_*l*_)) has a corresponding gene in *G*′. Thus, if there was ever a loss of a gene copy along any edge of the species tree, it would have to be compensated for by either a gene duplication event or am additive transfer event to ensure that all species descended from that edge still have a copy of the gene. Since the constructed reconciliation of *G*′ and *S* does not have any gene duplications and the only additive transfer does not affect edges of *S*_*r*_ or *S*_*l*_, there can not be any losses in the constructed reconciliation. □

The following lemma establishes the forward direction of claim 1.

##### Lemma 3.9.

*If there exists a YES answer to the D-rSPR problem on ρ then there exists a YES answer to the D-DTRL instance δ with reconciliation cost at most* 10*n* + 5*k* − 4.

*Proof.* Lemma 3.7 shows that the constructed reconciliation of *G*′ an *S* is a valid DTRL-reconciliation, and Lemmas 3.7 and 3.8 imply that this reconciliation does not have any losses or duplications. Furthermore, if |𝒫| ≤ *k* then, based on our construction and on Observation 1, |𝒫′| ≤ *k*. Thus, the constructed reconciliation of *G*′ and *S* has at most 2*n* + *k* − 2 nodes that represent replacing transfers, with at most *k* replacing transfers corresponding to the nodes of 𝒫′ and exactly 2*n* − 2 replacing transfers corresponding to the set *X*_*G*_. Finally, *rt*(*G*) represents an additive transfer event. Thus, the reconciliation cost of *G*′ and *S* is at most 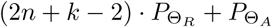 which is 10*n* + 5*k* − 4. □

### 3.4 Proof of Claim 1: Reverse direction

Conversely, we now assume that we have a YES answer to the D-DTRL instance *δ* with reconciliation cost at most 10*n*+ 5*k* − 4, and will show that there must then exist a solution of size at most *k* to the D-rSPR instance *ρ*. In this proof, we will first characterize the structure of any optimal DTRL-reconciliation of *G* and *S*, and then show that this structure implies the existence of a specific set of evolutionary events.

The next three lemmas identify basic properties of any optimal DTRL-reconciliation of *G* and *S* and follow easily based on the construction of the gadget. Specifically, the first lemma follows directly from the close correspondence between the topologies of *G*_*l*_ and *S*_*r*_, the second lemma follows from the presence of the 10 extraneous leaves on the path from *rt*(*S*) to *S*_*r*_, and the third lemma follows easily from the specific construction of the nodes in *Y*_*G*_ in the gene tree gadget.

#### Lemma 3.10.

*Given any optimal DTRL-reconciliation for G and S, any internal node g* ∈ *I*(*G*_*l*_) *must map to lca*_*S*_(ℒ(*G*(*g*))) *and represent a speciation event.*

#### Lemma 3.11.

*Given any optimal DTRL-reconciliation for G and S, no node of G maps to rt*(*S*).

#### Lemma 3.12.

*Given any optimal DTRL-reconciliation for G and S, each node y* ∈ *Y*_*G*_ *must represent a replacing transfer event.*

The next lemma shows that in any optimal DTRL-reconciliation of *G* and *S*, the number of gene copies present in any node (or edge) or the species tree is at most 1. The idea behind the proof of this next lemma is illustrated in Figure 4.

**Fig. 4.**
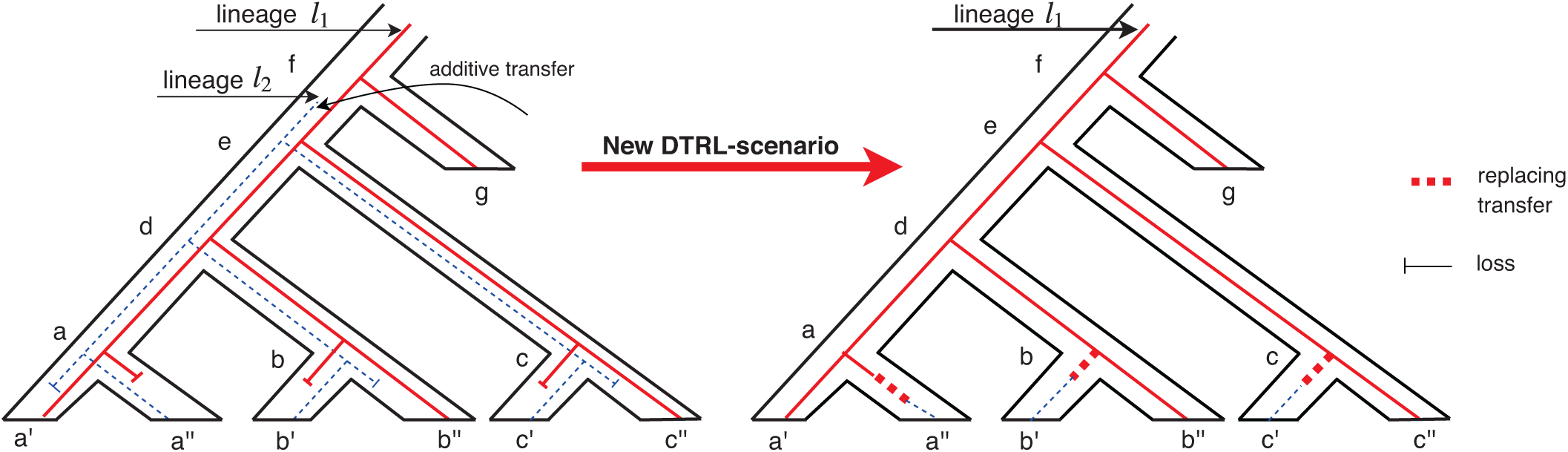
A illustration of the idea behind the proof of Lemma 3.13. The figure on the left shows the evolution of two gene lineages, lineage *l*_1_ (solid red lines) and lineage *l*_2_ (dashed blue lines), inside the species tree (the tube tree). According to that evolutionary scenario, both lineages are present in the species *a, b, c, d*, and *e* of the species tree. Observe that the corresponding DTRL-reconciliation would invoke six losses, in addition to the additive transfer that created lineage *l*_2_. The figure on the right shows how this evolutionary scenario can be modified by removing lineage *l*_2_ and instead invoking replacing transfer events to replace the genes from lineage *l*_1_ in nodes *a*′′, *b*′, and *c*′. Observe that the corresponding DTRL-reconciliation invokes six fewer losses and one fewer additive transfer but has 3 more replacing transfers. Thus, based on our assigned event costs, the DTRL-reconciliation corresponding to the evolutionary scenario on the left could not have been optimal.

#### Lemma 3.13.

*Given any optimal DTRL-reconciliation for G and S, there does not exist any node of S with more than one gene copy.*

*Proof.* Suppose for contradiction that at least one such node *a* ∈ *V*(*S*) exists. Without loss of generality we can assume that *a* is the first such node in a post-order traversal of *S*. Since, each leaf node of *S* has at most one gene copy, *a* must be an internal node. Thus, let *a*′ and *a*″ denote the two children of *a*. By our assumption, both *a*′ and *a*″ have at most one gene copy, while *a* has at least 2. Thus, there must be at least one loss along the edge (*a, a*′) and at least one loss along the edge (*a, a*″). We will show how to modify this current DTRL-reconciliation and reduce the total reconciliation cost. For simplicity, we will assume that *a* has exactly two gene copies, but the proof easily generalizes to greater than two gene copies.

We will modify the current DTRL-reconciliation as follows: Instead of incurring one loss at each of the two children edges of *a*, we move this loss upwards to the edge (*pa*(*a*), *a*), reducing the number of gene copies in *a* by 1. At least one of *a*′ or *a*″, say *a*′. must have inherited its single gene copy from the surviving gene lineage. Thus, the gene lineage entering *a*′ would be unaffected by the loss of the other copy in *a*. The other child *a*″ may have received its copy from the deleted lineage, and so may be affected by the loss at *a*. This can be resolved by invoking a replacing transfer event to replace the gene lineage coming into *a*″ from *a* with the desired gene lineage. Such a replacing transfer can always be added, if it does not already exist, at the parent node of the node from *G* that maps to *a*″ (or to its closest descendant if no node maps directly to *a*″).

We apply this modification iteratively towards the root of *S*, until the first (or highest) node along this path with the additional gene copy is reached. The source of the additional gene copy at this node must be either a gene duplication or additive transfer event on the gene tree. By removing the extra gene copy at this node, we therefore also reduce the number of gene duplications or additive gene transfers by 1. Overall, during this iterative process, we reduce the number of losses at each iteration by 2, add at most one replacing transfer event per iteration, and replace at least one duplication or additive transfer event by a speciation event during the last iteration. Based on our assigned event costs, this results in a net reduction in the total reconciliation cost. Since the initial DTRL-reconciliation of *G* and *S* was optimal, this is a contradiction. Thus, there cannot be any nodes in *S* with more than one gene copy. □

The following corollary follows immediately based on the proof of the previous lemma.

#### Corollary 3.1.

*Given any optimal DTRL-reconciliation for G and S, there does not exist any node in G that represents a duplication event.*

#### Lemma 3.14.

*There are no gene losses in any optimal DTRL-reconciliation for G and S.*

*Proof.* By Lemma 3.13 we know that each node of *S* has at most one gene copy. We also know that each leaf node of the species tree node *a* ∈ *Le*(*S*(*S*_*r*_)) ∪ *Le*(*S*(*S*_*l*_)) has a corresponding gene in *G*′. Thus, if there was ever a loss of a gene copy along any edge of the species tree, it would have to be compensated for by either a gene duplication event or an additive transfer event to ensure that all species descended from that edge still have a copy of the gene. By Corollary 3.1 we know that *G* does not have any duplication nodes in any optimal DTRL-reconciliation. Furthermore, since any node of *S* has at most one gene copy (Lemma 3.13), any additive transfer event not the root of *G* would either be preceded by a gene loss in the recipient lineage or would be immediately followed by a gene loss so as not to have more than one gene copy in any node of *S*. Thus, it would be possible to substitute any such additive transfer with a replacing transfer event and reducing the number of gene losses. However, this would lead to a DTRL-reconciliation with lower reconciliation cost, a contradiction. Thus, since the only additive transfer may occur at the root of the gene tree, and there are no gene duplications, there cannot be any gene losses in any optimal DTRL-reconciliation of *G* and *S*. □

#### Lemma 3.15.

*Given any optimal DTRL-reconciliation for G and S, then there is exactly one node that represent additive transfer.*

*Proof.* By the proof of Lemma 3.14 above, we know that the only possible additive transfer node is *rt*(*G*). It therefore suffices to prove that *G* must have at least one additive transfer event. By Lemma 3.11 we know that no node of *G* maps to *rt*(*S*), and by Lemma 3.10 we know that node *rt*(*G*_*l*_) maps to a node of *V*(*S*_*r*_). Without an additive transfer event bringing a copy of the gene to nodes of *S*_*l*_, the number of gene copies in nodes of *S*_*l*_ would be zero, a contradiction. □

The following lemma establishes the reverse direction of claim 1.

#### Lemma 3.16.

*Given any optimal DTRL-reconciliation for G and S with cost at most* 10*n* + 5*k* − 4, *there exists a solution for the D-rSPR instance ρ of size at most k.*

*Proof.* Based on Lemmas 3.14 and 3.15 and Corollary 3.1, we know that any optimal DTRL-reconciliation of *G* and *S* must invoke exactly one additive transfer, no duplications, and no losses. Thus, since the total reconciliation cost is at most 10*n* + 5*k* − 4, the total number of replacing transfers can be no more than 2*n* − 2 + *k*. Now, by Lemma 3.12 we know that each of the 2*n* − 2 nodes in *Y*_*G*_ must be replacing transfers. Thus, the number of nodes of *Z*_*G*_ that are replacing transfers is at most *k*, and the number of nodes of *Z*_*G*_ that represent speciation events is at least *n* − 1 − *k* (since |*Z*_*G*_| = *n* − 1).

Observe that, according to our gadget, the original tree *T* from the D-rSPR instance *ρ* corresponds to subtree *G*_*r*_ of the gene tree and tree *T*′ corresponds to subtree *S*_*l*_ of the species tree. Also observe that if a node from *Z*_*G*_ represents a speciation event then it must map to a node from *S*_*l*_. Therefore, there exist at most *k* internal nodes of *S*_*l*_ that are recipients of replacing transfer events (since *S*_*l*_ has exactly *n* − 1 internal nodes). Note that the corresponding transfer events on *G* must all be from *G*_*r*_, and let *A* denote the set of these corresponding transfer nodes from *G*_*r*_.

Now, consider the forest *F*_*S*_ created from *S*_*l*_ by cutting all edges that connect the at most *k* nodes that are recipients of replacing transfer events to the rest of *S*_*l*_. Likewise, consider the forest *F*_*G*_ created from *G*_*r*_ by first removing all nodes from *X*_*G*_ and collapsing all nodes with only one child (i.e., all nodes of *Y*_*G*_ are collapsed), and then cutting all edges that connect the nodes of *A* to the rest of the tree. It is not hard to argue that the two forests *F*_*S*_ and *F*_*G*_ must be identical, which provides a solution of size at most *k* for the D-rSPR problem on *T* and *T*′. □

## 4 Experimental Analysis

There do not currently exist any algorithms or heuristics to compute DTRL reconciliations, and it is not even known how algorithms for computing optimal DTL reconciliations perform when confronted with gene trees that have been affected by both additive and replacing transfers. Therefore, we first focused on answering two fundamental questions: (i) How is the accuracy of DTL reconciliation affected by the presence of replacing horizontal gene transfers? (ii) How well does DTL reconciliation perform at inferring replacing transfer events?

To answer these questions, we used the recently developed simulation framework SaGePhy [22] to stochastically evolve gene trees inside a given species tree under a model that allows for gene duplications, additive transfers, replacing transfers, and gene losses. Using this simulation framework we created a large number of gene trees with varying rates of evolutionary events, computed optimal DTL reconciliations for the gene/species tree pairs, and evaluated the accuracy of the inferred reconciliations by comparing them to the true evolutionary histories of those gene trees. To compute optimal DTL reconciliations we employed the widely-used RANGER-DTL [2], [3] software package.

### Simulated datasets

We used SaGePhy [22] to generate 100 species trees, each containing 100 leaves and of height 1, under a birth-death process. Next, inside each of the species trees, we generated three different gene trees using low, medium, and high rates of duplication, additive transfer, replacing transfer, and loss events, resulting in three sets of 100 gene trees. To generate the low DTRL gene trees, we used duplication, additive transfer, replacing transfer, and loss rates of 0.133, 0.133, 0.133, and 0.266, respectively; for the medium DTRL gene trees we used rates of 0.3, 0.3, 0.3, and 0.6, respectively; and for the high DTRL gene trees we used rates of 0.6, 0.6, 0.6, and 1.2, respectively. Thus, the total transfer rate was twice the duplication rate, with an equal rate of additive and replacing transfers, and the loss rate was assigned to be equal to the sum of the duplication and additive transfer rates. These duplication, transfer, and loss rates are based on rates observed in real data and capture both datasets with lower rates of these events and datasets with a very high rate of these events [4].

For the low DTRL gene trees, the average gene tree leaf set size was 96.11, with an average of 2.37 additive transfers, 2.65 replacing transfers, and 2.19 duplication events per gene tree. For the medium DTRL gene trees, the average gene tree leaf set size was 94.75, with an average of 5.09 additive transfers, 5.01 replacing transfers, and 5.00 duplication events per gene tree. For the high DTRL gene trees, the average gene tree leaf set size was 110.22, with an average of 9.52 additive transfer events, 9.42 replacing transfer events, and 10.39 duplication events per gene tree.

#### 4.1 Impact of replacing transfers on DTL reconciliation

We evaluated the accuracy of DTL reconciliation in inferring the evolutionary event and species tree mapping for each internal node in the simulated gene trees. We computed a single optimal reconciliation for each gene tree using RANGER-DTL 2.0 [3] with default parameters (i.e., transfer cost of 3, duplication cost of 2, and loss cost of 1) and compared the computed reconciliation against the true evolutionary history of that gene tree. We observed very high accuracy for inferring the correct event type (speciation, duplication, or transfer) at each gene tree node. For instance, for the low DTRL gene trees, 99.67%, 96.35% and 96.22% of the gene tree nodes labeled as speciation, duplication, and transfer, respectively, in the computed reconciliations were inferred correctly. Even for the high DTRL gene trees, these percentages remained very high at 95.69%, 87.49%, and 95.25%, respectively. These results are shown in Figure 5(a).

**Fig. 5.**
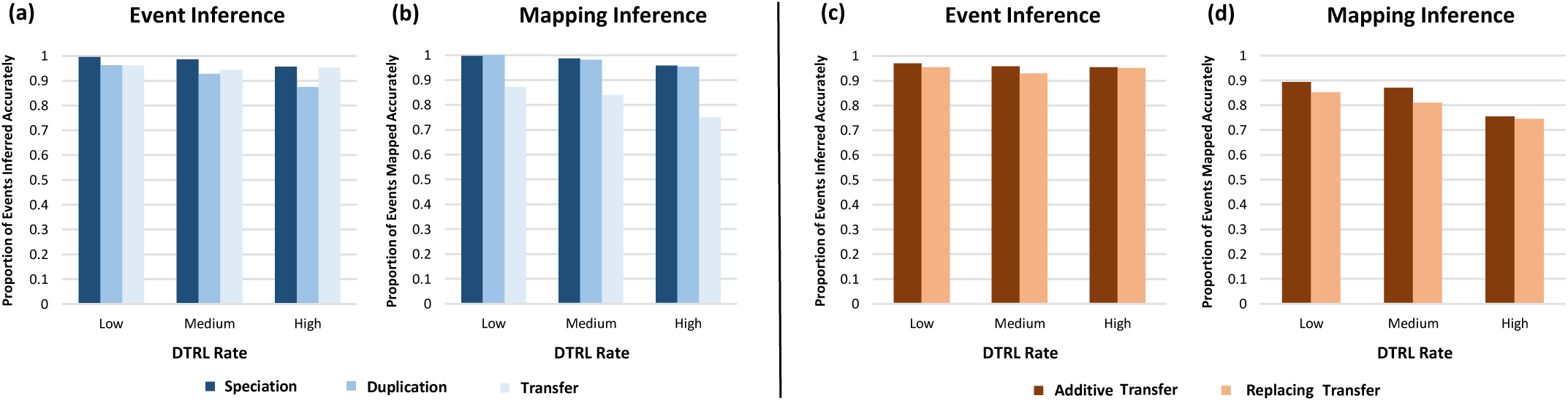
Accuracy of DTL reconciliation in the presence of replacing transfers. Part (a) shows the fraction of internal nodes across all low DTRL, medium, DTRL, and high DTRL gene trees, whose event types, speciation, duplication, or transfer, are inferred correctly through DTL reconciliation. Part (b) shows the corresponding fractions for correct mapping inference. Part (c) shows the fraction of additive transfer nodes and replacing transfer nodes across all low DTRL, medium, DTRL, and high DTRL gene trees, that are correctly inferred as transfer events by DTL reconciliation. Part (d) shows the corresponding fractions for correct mapping inference. For each DTRL rate, results are averaged across 100 datasets.

Looking at the accuracy of mapping inference, we found that 99.09%, 97.11%, and 92.15% of all internal nodes were assigned the correct species node mapping for the low, medium, and high DTRL gene trees, respectively. Detailed results are shown in Figure 5(b).

We compared these results for event and mapping accuracy with results obtained on gene trees simulated with the same overall rates of duplication, transfer, and loss events but in which all simulated transfers were additive transfers (no replacing transfers). We found that the numbers were nearly identical, showing that the presence of replacing transfers does not negatively affect the accuracy of DTL reconciliation itself. For example, for the high DTL gene trees, the percentage of speciation, duplication, and transfer nodes assigned the correct event type was 95%, 81%, and 95%, respectively, and 91% of all nodes were assigned the correct mapping. Note, however, that DTL reconciliation cannot distinguish between additive and replacing transfers, and both types of transfer events are simply inferred as “transfers”.

### Accuracy of inferring replacing transfers

Next, we performed additional analysis to study if there was any discrepancy in the accuracies of inferring the correct event type (transfer) or mapping for additive transfers and those for replacing transfers. For the low DTRL gene trees, we found that additive transfers were assigned the correct event type 97.05% of the time and the correct mapping 89.45% of the time, while for replacing transfers these numbers were 95.47% and 85.28%, respectively. Likewise, for the medium DTRL gene trees, additive transfers were assigned the correct event type 95.87% of the time and the correct mapping 87.03% of the time, while for replacing transfers these numbers were 93.01% and 81.04%, respectively. For high DTRL gene trees, these numbers were 95.38% and 75.53% for the additive transfers and 95.12% and 74.52% for the replacing transfers. Overall, this shows that replacing transfers are inferred and mapped with accuracy comparable to that of additive transfers. These results are shown in Parts (c) and (d) of Figure 5.

These results are highly significant and suggest that, to design an effective heuristic for DTRL reconciliation, it may suffice to first use DTL reconciliation to identify transfer events and then classify those transfer events as being either replacing or additive.

## 5 A Heuristic for Classifying Transfers

To explore the feasibility of accurately classifying transfer events inferred through DTL reconciliation, we designed a simple heuristic for classifying inferred transfers and tested its accuracy on several simulated datasets. Given gene tree *G* and species tree *S*, our heuristic first computes an optimal DTL reconciliation, initially classifies all inferred transfer events as additive, and then greedily attempts to reclassify some of these transfer events as replacing. To determine if a transfer can be replacing, the heuristic checks if the resulting loss of a gene lineage in the recipient species will make it impossible to generate at least as many gene copies at each leaf descendant of the recipient species as are actually present. If it does then the transfer remains an additive transfer but otherwise is reclassified as a replacing transfer.

This heuristic thus depends only on the actual counts (based on *G* and *S*) and implied/inferred counts (based on the computed reconciliation) of genes at each leaf of the species tree.

More precisely, the heuristic works as follows:

1. Calculate the number of gene copies from *Le*(*G*) that are present in each extant species represented in the species tree. For a species *s* ∈ *Le*(*S*), this count is represented by *actual-count*(*s*). Note that this is simply the number of leaf nodes of *G* that map to leaf *s*.
2. Compute an optimal DTL reconciliation for *G* and *S* (using RANGER-DTL 2.0 [3] with default parameters).
3. Classify each inferred transfer event as an additive transfer.
4. Based on the current reconciliation, compute the number of gene copies that would occur in each extant species if there were no gene losses. This can be counted easily as follows: Consider the path from the root of the species tree to the species (leaf) under consideration. Count the number of gene duplication nodes on the gene tree that map to a node on this path; let this number be denoted *n*_1_. Count the number of additive transfer events on the gene tree whose recipient is a node on this path; let this number be denoted *n*_2_. Determine if the root of the gene tree maps to a node on this path; If so, assign *n*_3_ = 1, otherwise *n*_3_ = 0. The final required count is simply *n*_1_ +*n*_2_ +*n*_3_. For a species *s* ∈ *Le*(*S*), this final count is represented by *inferred-count*(*s*).
5. For each node *g* in a pre-order traversal of *G*:
  a. If *g* is a transfer event:
    i. Let *x* ∈ *V*(*S*) denote the recipient species for that transfer event.
    ii. Check if *inferred-count*(*s*) > *actual-count*(*s*) for each *s* ∈ *Le*(*S*(*x*)). If yes, reclassify *g* as a replacing transfer and reduce *inferred-count*(*s*) by 1 for each *s* ∈ *Le*(*S*(*x*)).
6. Output the resulting classification of inferred transfer events.

An implementation of the heuristic algorithm is freely available open-source as part of the RANGER-DTL software package: https://compbio.engr.uconn.edu/software/ranger-dtl/. Next, we illustrate this algorithm through an example.

### Illustration of the heuristic

Consider the gene tree and species tree shown in Figure 6. Suppose the inferred reconciliation of the gene tree and species tree labels gene node *g*_3_ as a transfer event mapping to species node *s*_4_ and with recipient species *s*_6_, gene node *g*_6_ as a gene duplication mapping to species *D*, gene node *g*_7_ as a transfer event mapping to species node *G* and with recipient species *A*, and all other nodes as speciations with the root of the gene tree mapping to the root of the species tree. The heuristic starts with this reconciliation and its task is to assign each of the two transfer nodes at *g*_3_ and *g*_7_ to be additive or replacing.

**Fig. 6.**
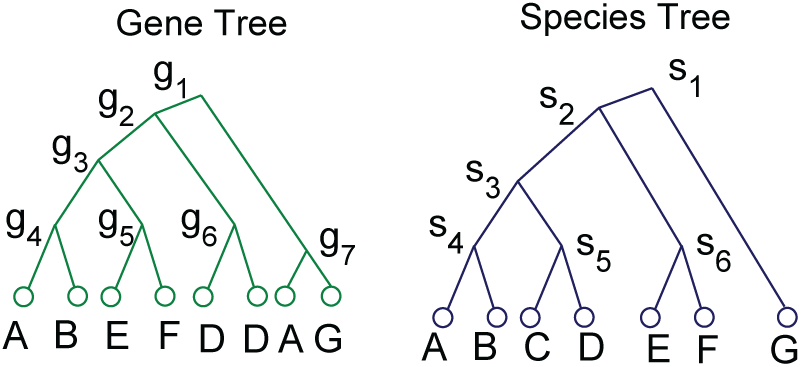
Gene tree and species tree for illustrating heuristic.

The *actual-counts* at the leaves *A, B, C, D, E, F*, and *G* of the species tree are easily computed to be, respectively: 2, 1, 0, 2, 1, 1, 1.

The heuristic starts by assuming that both transfer events are additive and that there are no losses, and computes the initial values of the *inferred-counts* at leaves *A, B, C, D, E, F*, and *G* of the species tree, respectively, as follows: 2, 1, 1, 2, 2, 2, 1.

Note that each of these *inferred-counts* is at least equal to the corresponding *actual-count*.

Next, the heuristic considers all transfer nodes on the gene tree one at a time in pre-order. Thus it first considers node *g*_3_. It checks to see if the additive transfer at *g*_3_ could have been a replacing transfer instead. To make this determination, it checks if the inferred-count at each leaf descended from the recipient of this transfer event (i.e., all leaves descended from node *s*_6_) is strictly greater than its corresponding *actual-count*. In this example, we would check to make sure that *inferred-count*(*E*) > *actual-count*(*E*) and *inferred-count*(*F*) > *actual-count*(*F*). Both inequalities hold true in this case, and so the transfer at node *g*_3_ is labeled as a replacing transfer. The next step is to update the *inferred-counts* to account for this change. The new *inferred-counts* at the leaves *A, B, C, D, E, F*, and *G* of the species tree, respectively, are thus: 2, 1, 1, 2, 1, 1, 1.

Continuing the pre-order traversal the heuristic then considers the transfer at node *g*_7_, and again checks to see if this additive transfer event could have been a replacing transfer. To make this determination, it checks if *inferred-count*(*A*) > *actual-count*(*A*). This inequality does not hold since both counts are 2. Thus, the heuristic labels the transfer at *g*_7_ as an additive transfer.

#### 5.1 Experimental Results

To evaluate the ability of this heuristic to classify transfer events accurately we applied it to several simulated datasets covering a wide range of evolutionary scenarios. We divide these datasets into three groups: Group 1 consists of the datasets described in Section 4 (consisting of the three sets of low, medium, and high DTRL trees) and we refer to these as *mixed datasets* since gene trees in these datasets contain both additive and replacing transfers. Group 2 consists of datasets in which all transfers are additive transfers. As before, this group is composed of three sets of low, medium, and high DTRL trees. Finally, group 3 consists of datasets in which all transfers are replacing transfers, divided as before into three sets of low, medium, and high DTRL trees. For group 2 and group 3 datasets, the duplication, transfer, and loss rates used to generate the low, medium, and high DTRL trees are identical to those used for group 1 (described in detail in Section 4), except that in group 2 all transfers are additive and in group 3 all transfers are replacing. Thus, the total number of duplication and transfer events are roughly the same across the three groups.

To evaluate the classification accuracy of our heuristic, we measured the following for each dataset from each of the three groups: (1) What fraction of all transfer events in the true evolutionary history of a gene tree are correctly inferred as transfer events by the heuristic. (2) What fraction of additive transfer events in the true evolutionary history of a gene tree are correctly inferred as additive transfers by the heuristic. And (3) what fraction of all replacing transfer events in the true evolutionary history of a gene tree are correctly inferred as replacing transfers by the heuristic. Table 1 shows these results. As also seen in Section 4, transfer events can be identified with high accuracy across all three groups and all three DTRL rates. Results are more variable for classification of the inferred transfers as being either additive or replacing (which is the primary task of the heuristic). In general, over 80% of the additive transfers and 86% of the replacing transfers are classified correctly for the low DTRL datasets across the three groups, 60%– 65% of additive and 80%–88% of replacing transfers are classified correctly for the medium DTRL datasets across the three groups, and approximately 52% of additive and 70% of replacing transfers classified correctly for the high DTRL datasets. These results show that the proposed heuristic is quite accurate at classification when DTRL rates are low but performance suffers as the rates increase. These results also show that, in general, the heuristic infers replacing transfers with greater accuracy than additive transfers. This is not entirely surprising given that the heuristic attempts to label as many of the transfers as replacing as possible; in particular, additive transfers may appear to be replacing due to the high loss rate in our datasets. As a result, the false negative rate for replacing transfers is low but the false positive rate is high (since many additive transfers may be classified as replacing), while the opposite is true for additive transfers.

**TABLE 1.**
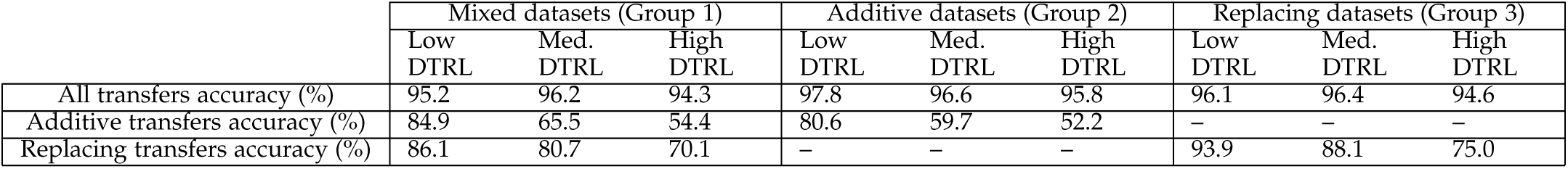
Classification accuracy of the heuristic. This table shows the results of applying the heuristic algorithm to the three groups of simulated datasets; Group 1 where gene trees contain both additive and replacing transfers, Group 2 where gene tree contain only additive transfers, and Group 3 where gene trees contain only replacing transfers. Each group is further divided into low, medium, and high DTRL datasets. For each dataset in each group, we measure (i) the percentage of true transfers that are inferred as transfers by the heuristic (“All transfers accuracy”), (ii) the percentage of true additive transfers that are inferred as additive transfers by the heuristic (“Additive transfers accuracy”), and (iii) the percentage of true replacing transfers that are inferred as replacing transfers by the heuristic (“Replacing transfers accuracy”). All numbers are averaged over the 100 gene tree/species tree pairs in each dataset. Note that the numbers reported for “All transfers accuracy” for Group 1 datasets are slightly different from those reported in Section 4. This is because there often exist multiple optimal reconciliations and each run of RANGER-DTL or the heuristic samples one of these optimal reconciliations at random.

|  | Mixed datasets (Group 1) |  |  | Additive datasets (Group 2) |  |  | Replacing datasets (Group 3) |  |  |
| --- | --- | --- | --- | --- | --- | --- | --- | --- | --- |
|  | Low<br>DTRL | Med.<br>DTRL | High<br>DTRL | Low<br>DTRL | Med.<br>DTRL | High<br>DTRL | Low<br>DTRL | Med.<br>DTRL | High<br>DTRL |
| All transfers accuracy (%) | 95.2 | 96.2 | 94.3 | 97.8 | 96.6 | 95.8 | 96.1 | 96.4 | 94.6 |
| Additive transfers accuracy (%) | 84.9 | 65.5 | 54.4 | 80.6 | 59.7 | 52.2 | – | – | – |
| Replacing transfers accuracy (%) | 86.1 | 80.7 | 70.1 | – | – | – | 93.9 | 88.1 | 75.0 |

Overall, these experimental results demonstrate that there is often sufficient information in DTL reconciliations to be able to distinguish between additive and replacing transfers, and suggest that classification of transfer events inferred through DTL reconciliation is a promising approach for estimating optimal DTRL reconciliations. While our current heuristic is simple and has limited classification accuracy, our experimental results do also suggest that, in general, the ability to distinguish between replacing and additive transfers based purely on gene tree (and species tree) topology diminishes rapidly as the rate of evolutionary events increases. This is not surprising since evolutionary events that occur after an additive or replacing transfer can completely erase the phylogenetic (i.e., topological) signature of that additive or replacing transfer. Nonetheless, we expect more advanced heuristics to be more effective at distinguishing between additive and replacing transfers even for high rates of evolutionary events.

## 6 Conclusion

Accurate detection of both replacing and additive transfer events is crucial for understanding horizontal gene transfer in microbes and understanding microbial evolution in general. In this work, we address this problem by formalizing and experimentally studying the DTRL reconciliation framework that simultaneously models gene duplication, loss, and both additive and replacing transfer. Our framework builds upon the traditional DTL reconciliation model and extends it substantially to properly model replacing transfers. We prove that the underlying computational problem is NP-hard, and our proof establishes a close relationship between the rSPR distance problem and DTRL reconciliation. Our experimental results show that DTL reconciliation, which assumes all transfers are additive, is surprisingly robust to the presence of replacing transfer, and suggest that it should be possible to design effective heuristics for the DTRL reconciliation problem based on DTL reconciliation. To explore the feasibility of such an approach, we devised a simple heuristic to classify inferred transfer events as being either additive or replacing and found that it achieves fairly good classification accuracy for low and medium rates of evolutionary events. This demonstrates the feasibility of estimating optimal DTRL reconciliations based on optimal DTL reconciliations followed by classification of inferred transfer events, and we expect improved heuristics to achieve greater classification accuracy. Our current heuristic has several limitations, of which the following two are particularly notable: First, it does not directly solve the DTRL reconciliation problem and its accuracy is therefore limited by the accuracy of the inferred DTL reconciliation (which does not model hidden events). Second, it ignores the presence of multiple optimal DTRL (or even DTL) reconciliations. Addressing these limitations may yield improved heuristics for DTRL reconciliation.

Our experimental results also suggest that, as expected, the ability to distinguish between replacing and additive transfers based purely on phylogenetic incongruence diminishes rapidly as the rate of evolutionary events increases, and therefore alternative approaches may be needed for such cases. One such alternative approach for estimating optimal DTRL reconciliations is to make use of available gene order information for the extant species in the analysis to classify each transfer event inferred through DTL reconciliation as being either additive or replacing based on genomic context. However, the applicability of such an approach is limited since it requires the use of complete genomic information and, due to genome rearrangements, can only be used for relatively closely related sets of species. A hybrid approach that uses both gene ordering information and phylogenetic incongruence may help overcome the limitations of the two separate approaches, and developing this hybrid approach is a promising research direction.

Finally, it would be useful to develop exact algorithms for the DTRL reconciliation problem. Even though we showed the problem to be NP-hard, it may be possible to design fixed parameter algorithms that can be efficiently applied to gene trees with small reconciliation cost (see. e.g., [14]), or to design effective branch and bound algorithms to rapidly compute optimal DTRL reconciliations for small gene trees.

## Acknowledgements

The authors thank Abhijit Mondal for pointing out an error in an initial implementation of the heuristic.

## Footnotes

1. Note that the DTL reconciliation model [31], [2] on which our new model is based allows the inferred reconciliation to be time-inconsistent. This is simply because imposing time consistency makes the DTL reconciliation problem NP-hard [31], while allowing time-inconsistency makes the problem efficiently solvable with negligible impact on accuracy.

